# Plant species richness and the root economics space drive soil fungal communities

**DOI:** 10.1101/2024.03.20.585751

**Authors:** Justus Hennecke, Leonardo Bassi, Cynthia Albracht, Angelos Amyntas, Joana Bergmann, Nico Eisenhauer, Aaron Fox, Lea Heimbold, Anna Heintz-Buschart, Thomas W. Kuyper, Markus Lange, Yuri Pinheiro Alves de Souza, Akanksha Rai, Marcel Dominik Solbach, Liesje Mommer, Alexandra Weigelt

## Abstract

Trait-based approaches have been increasingly used to relate plants to soil microbial communities. However, the plant organs mediating this plant-microbe interaction – the roots – have been largely overlooked. The recent discovery of the root economics space offers a predictive framework for the structure of soil microbial communities, and specifically soil-borne fungal communities. Applying this novel approach, our study in a grassland plant diversity experiment reveals distinct root trait strategies at the level of the plant community. In addition to significant effects of plant species richness, we show that both axes of the root economics space – the collaboration and conservation gradient – are strong drivers of the composition of the different guilds of soil fungi, including saprotrophic, plant pathogenic, and mycorrhizal fungi. Our results illustrate that the root economics space and plant species richness jointly determine the effects of plants on fungal communities and their potential role in plant health and ecosystem functioning.

## Introduction

Soil fungi can act as mutualists or antagonists of plants and thus promote or weaken the functioning of plants and ecosystems^1–3^. Understanding the drivers of the guild composition of fungal communities is important for the understanding of ecological processes that shape plant communities and crucial for the management of soil microbial communities^4–6^. Plant communities can exert strong selective pressure on soil fungal communities^7,8^. Both plant species richness^8^ and composition^9,10^ and, therefore, the associated plant traits are important components of belowground plant-fungal relationships. In the last decades, trait-based approaches have emerged to give valuable mechanistic insights into plants as drivers of the soil microbial community^11,12^ but these largely lack a belowground trait perspective.

Root traits are not just analogs of leaf traits^13^; their variation along two orthogonal axes has been described in the so-called root economics space (RES)^11^. While roots, like leaves, vary in root tissue density and relative root nitrogen content along the ‘fast–slow’ axis of the conservation gradient, there is an additional trade-off in specific root length and mean root diameter^11^. This trade-off has been explained by the interaction between roots and their associated mutualistic arbuscular mycorrhizal fungi (AMF), with thicker roots generally hosting more AMF (‘outsourcing’ strategy) and thinner roots maximizing root surface for independent nutrient uptake (‘do-it-yourself’, ‘DIY’ strategy)^11,14^. Recent studies have largely confirmed this global trait coordination of species across regions and vegetation types^15–17^. However, it is unclear to what extent the species-level RES is also represented at the plant community-level^18,19^. Therefore, understanding how root functional strategies scale from the species-level to the community-level is critical to utilizing the RES as a trait-based framework for soil microbial communities and ecosystem functioning.

Multiple components of the fungal community can be affected by abiotic and biotic factors. Based on a functional classification, soil fungal communities are composed of three main guilds: saprotrophs, pathogenic, and mycorrhizal fungi^20^. While the overall fungal abundance (or biomass) can change e.g. through increased resource availability^21^, changes in the functional or taxonomic composition of the community likely result from interspecific processes such as competition or differences in resource utilization among taxa or guilds^22^. Approaches integrating quantitative and qualitative measures of fungal communities are needed for a holistic view of the effects of plants on fungal communities, especially as both components are relevant to ecosystem processes^23^. Generally, soil fungal communities are closely linked to the plant community, as they are dependent on carbon input from the plant but also drive the nutrient cycling and availability to the plant^24^. However, frameworks that explicitly link plant traits to the functional composition of soil fungal communities are limited in number and explanatory power^25,26^, and studies that empirically tested trait-based frameworks have largely omitted root traits^27^. The additional complexity of trait variation in roots compared to leaves is not yet integrated into such mechanistic frameworks despite its potential opportunities to reconcile trait-based approaches^28^ and better understand plant communities as drivers of soil fungal communities.

If the RES exists at the community-level, it can be used to extend the initial framework of Wardle *et al.*^12^ linking the ‘fast–slow’ plant trait gradient to the microbial community. Recently, Hennecke *et al.*^17^ presented a theoretical framework of how root trait gradients link with the composition of fungal communities in the rhizosphere. Plant communities with dominating traits on the ‘outsourcing’ end of the collaboration gradient should accommodate a higher diversity and relative abundance of AMF and, due to their protective role^29^, less plant pathogenic fungal diversity and relative abundance^30,31^. Further, plant pathogenic fungi should benefit more from higher nutrient availability and lower defense of plant tissue, both of which align with a ‘fast’ strategy of the growth-defense trade-off^32^ along the conservation gradient. Saprotrophic fungi strongly depend on the quality and quantity of available plant litter^12,33^.

Roots at the slow end of the conservation gradient produce low-quality litter, yet it is unclear whether this affects saprotroph diversity and abundance^17^.

In addition to plant functional traits, plant species richness can also cause differences in soil fungal composition^34,35^. Higher richness of primary producers can influence the composition and diversity of soil microbes via increased heterogeneity of resources, including roots, exudates, and litter^36–38^. Additionally, at similar soil fertility, plant species richness is often correlated with primary productivity^39^, thereby increasing the amount of plant-based resources for fungi and hence fungal biomass^8,38^. Multiple studies found fungal diversity to increase with plant species richness^40–42^, but opposing or no effects were also reported^43,44^, indicating that the plant species richness-fungal diversity relationship can depend on environmental conditions^43^, scale^45^, or differ between fungal guilds. Plant pathogenic fungi are predicted to be less abundant due to decreased host density with increased plant species richness^46^. We, therefore, expect that fungal guild composition differs across the plant species richness gradient, with a stronger increase in the abundance and diversity of saprotrophic and arbuscular mycorrhizal fungi compared to plant pathogens.

In a grassland biodiversity experiment, we aimed to disentangle the effects of root traits and plant species richness on saprotrophic, plant pathogenic, and arbuscular mycorrhizal fungi as the most relevant fungal guilds in grassland soils. We test three overarching hypotheses: (1) Root trait organization at the plant community-level mirrors the RES (i.e. the collaboration and conservation gradients) previously found at the species-level. (2) The diversity and relative abundance of soil fungal guilds are structured by the community RES (3) Plant species richness is linked to increased plant biomass and thereby increases fungal diversity and biomass, but not all fungal guilds benefit equally: we expect fungal mutualists and saprotrophs to benefit more from plant species richness than plant pathogens.

Overall, we aim to advance the potential of trait-based frameworks by integrating root functional strategies. As a first major step, we show that the root trait gradients at the community-level are strong determinants of the diversity and relative abundance of soil fungal guilds. While both, root traits and plant species richness, are correlated with root biomass, soil fungal biomass is only driven by species richness and not root traits. Our study illustrates that root traits and plant species richness affect different properties of soil fungal communities and therefore jointly mediate the effects of plants on fungal communities.

## Results and discussion

### Root traits at the community-level

To determine root traits at the community-level, we sampled roots from bulk soil without separation by plant species. The PCA of these root traits shows two clear axes that explain a cumulative 79.4% of the variation (Fig. 1). The trait organization closely resembles the root economics space (RES) found at the species-level across a large number of species and biomes in Bergmann *et al.*^11^. The first axis of the varimax-rotated PCA, explaining 42.2% of the variation in community root traits, represents the collaboration gradient of the RES ranging from high root diameter (‘outsourcing’ strategies) to high specific root length (‘do-it-yourself’, ‘DIY’). The second axis explained 37.2% and represents the conservation axis ranging from high root tissue density (‘slow’) to high root nitrogen (‘fast’). This is in line with Da *et al.*^19^ who found the community RES to be nearly identical to the species-level RES when using community weighted-mean root traits of woody species in a temperate forest. In contrast, Lachaise *et al.*^18^ identified the RES on community weighted-mean traits in observational grasslands with root nitrogen not following expected patterns. Taken together, we show for the first time that directly measured plant community root traits, rather than community traits calculated from species-specific traits, follow the same functional trade–offs as at the species-level. This finding suggests that even under the same abiotic conditions, different economic strategies of a community can be successful. It further demonstrates the robustness of the trait organization across a wide plant species richness gradient but also highlights the need to better understand the conditions that lead to deviations from the RES found in other studies.

**Fig. 1:**
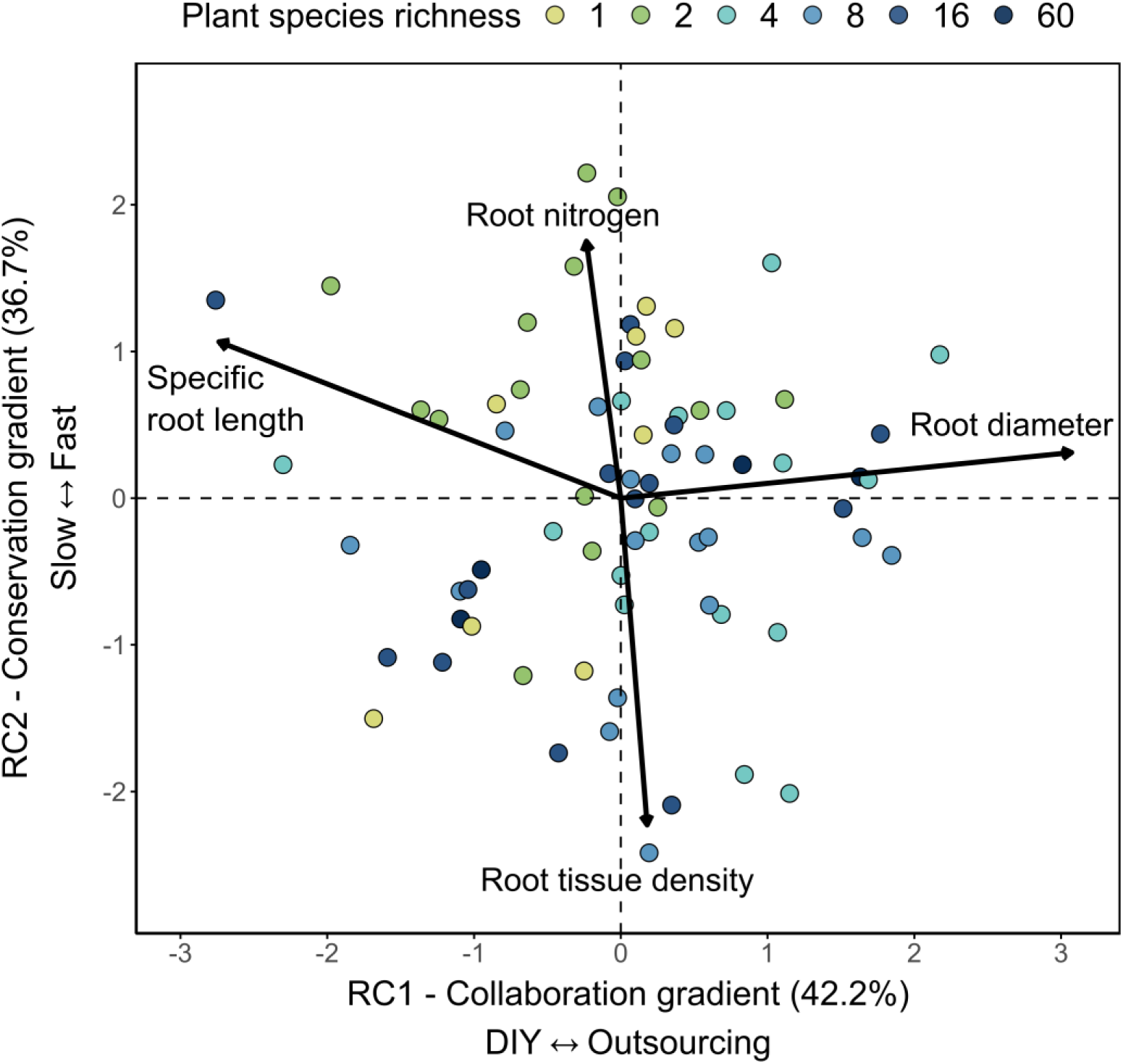
PCA of the community root traits. Each point represents a plant community (*n* = 73). The two axes closely resemble the species-level root economics space^11^. The traits of the collaboration gradient load on the first axis while traits of the conservation gradient load on the second axis. Varimax rotation was used to increase interpretability of the two axes. Points are color-coded for plant species richness of the plot. RC, rotated component.

The community root traits varied along the gradient of sown plant species richness. While the traits of the collaboration axis, represented by the scores of the first rotated component (RC1) of the PCA, were not significantly related to plant species richness (Estimate = 0.092, *P* = 0.404), scores along the conservation axis (RC2) showed a stronger relationship with plant species richness (Estimate = -0.246, *P* = 0.023, Fig. 2, Supplementary Table S1). Accordingly, more diverse plant communities were characterized by a ‘slower’, more resource-conservative strategy. This was primarily driven by the decrease of root nitrogen in more diverse plant communities (Supplementary Table S1), which, in line with other studies, can be attributed to a ‘nutrient dilution effect’ with higher plant biomass and increased nitrogen use efficiency^47,48^.

**Fig. 2:**
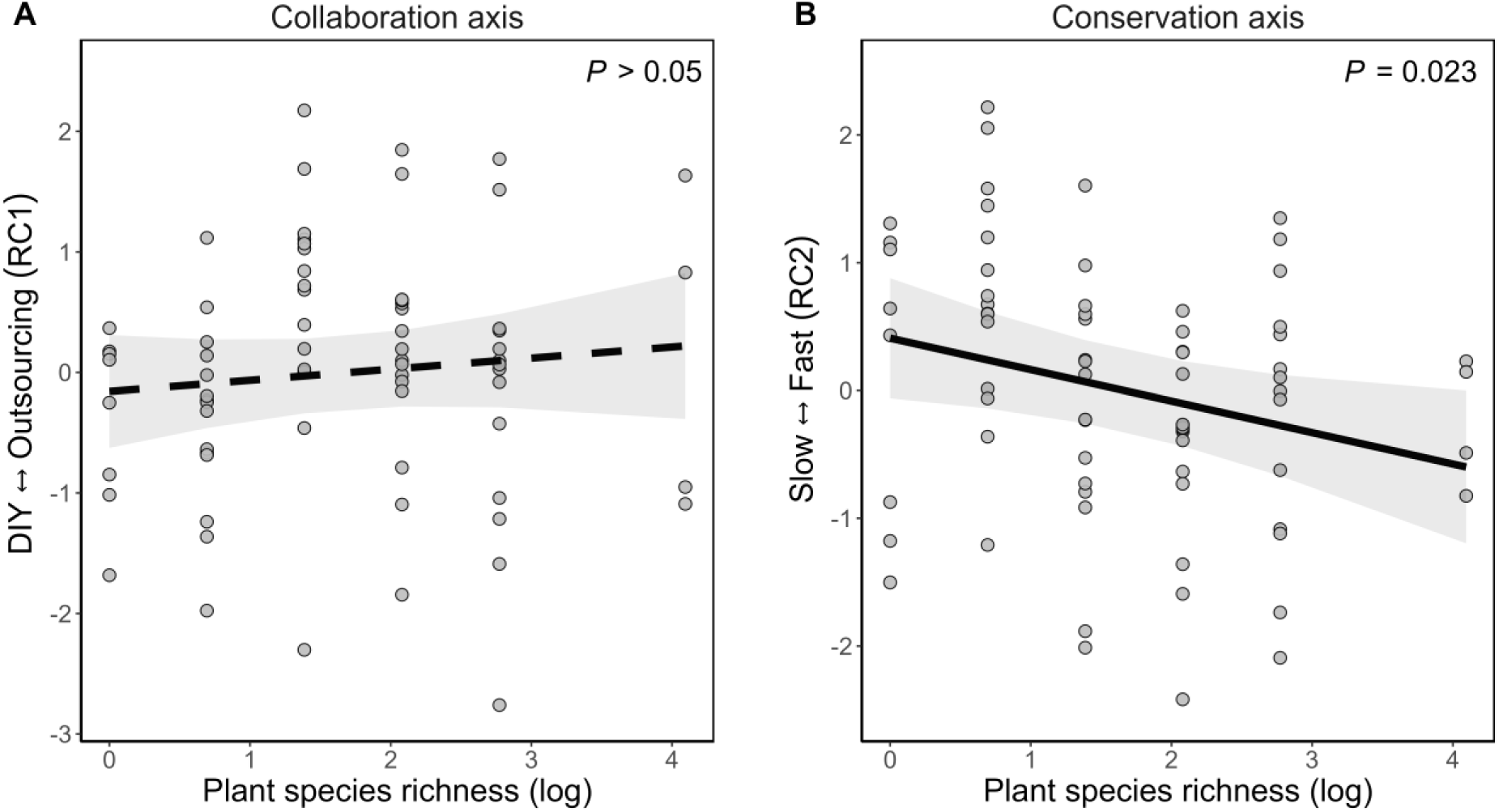
Change of root functional strategies along the plant species richness gradient. Scores of the first and second rotated component (RC) of the root trait PCA, representing the (A) collaboration axis and (B) conservation axis, in relation to the plant species richness gradient. Each point represents the plant community of one experimental plot (*n* = 73). Regression lines are based on mixed-effects model predictions, solid lines indicate significant relationships (P<0.05), dashed lines indicate non-significant relationships (P>0.05). The grey bands around the regression lines depict the 95% confidence interval.

### Links between root traits and soil fungal guilds

We analyzed how the diversity and relative abundance of saprotrophic, plant pathogenic, and arbuscular mycorrhizal fungi were related to the sown species richness and root traits of the plant communities. We found that the Shannon diversity of fungal saprotrophs was positively related to plant species richness (Table 1, Fig. 3), in line with our expectations. This suggests that plant species richness effects increase root biomass^49,50^ and reduce litter quality^51^, ultimately affecting saprotrophic diversity. In addition to plant species richness, the root functional strategies of the plant community had strong effects on the saprotrophic community. Easily-available carbon from high-quality litter in roots with ‘fast’ traits and exudates is also used by bacteria^12,52^ and therefore putatively less available to fungal saprotrophs, resulting in lower fungal saprotroph diversity (Table 1, Fig. 3). However, we found no significant change in saprotroph relative abundance with ‘fast’ root traits (Table 1, Fig. 3), suggesting that higher resource quality favors fewer fungal taxa that still form a similar proportion of the fungal community. ‘Outsourcing’ root strategies along the collaboration axis did not affect fungal saprotrophic diversity but significantly increased the relative abundance of saprotrophic fungi (Table 1, Fig. 3). As the mechanisms behind this are not obvious and previously reported relationships between the collaboration gradient of the RES and the saprotrophic fungal community and decomposition rates are variable^17^, we see this as an exciting avenue for further studies.

**Fig. 3:**
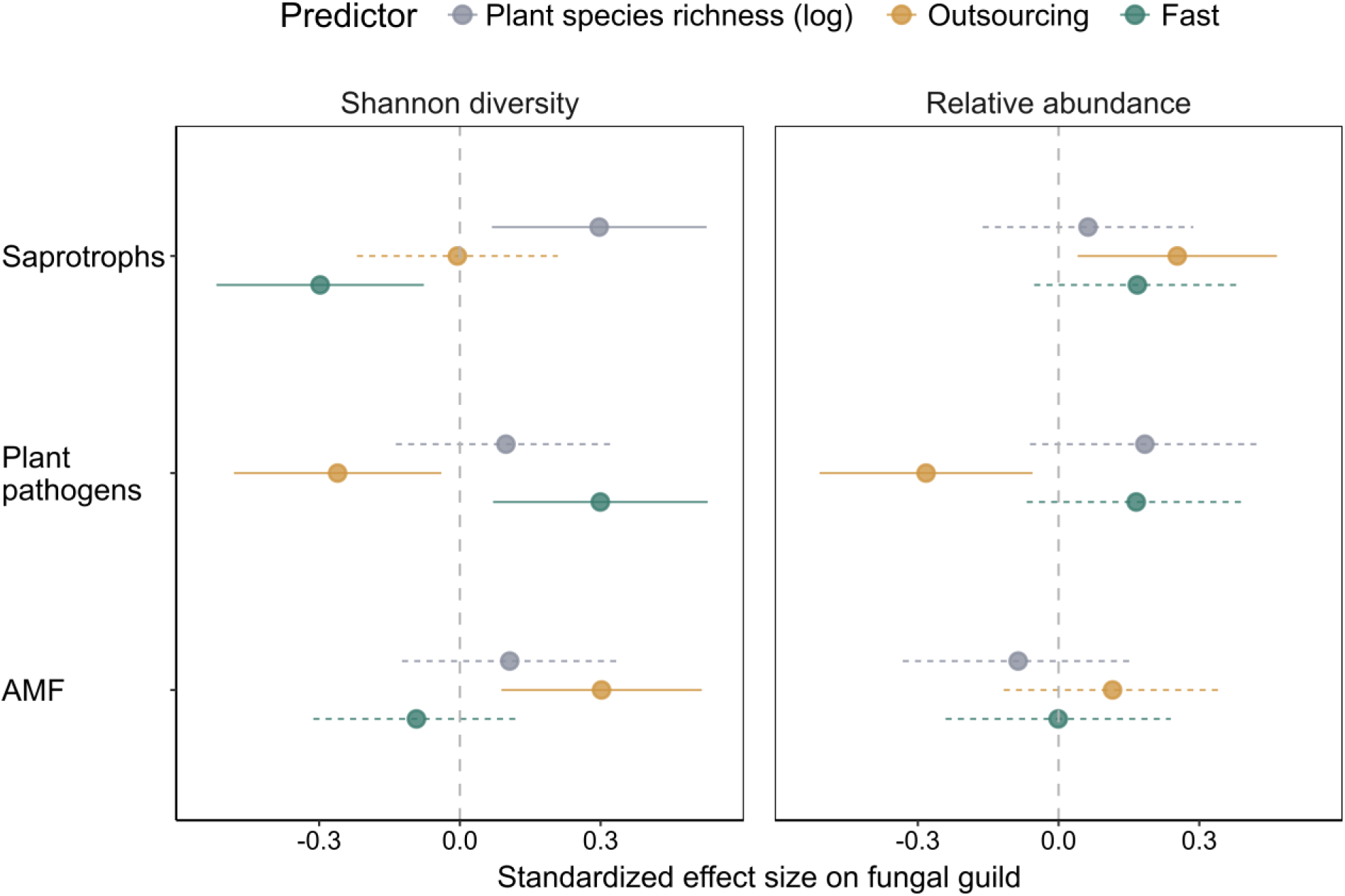
Effects of plant species richness and root trait gradients on fungal guilds. Standardized effect sizes of plant species richness (log) and root trait strategies (‘outsourcing’ along the collaboration gradient and ‘fast’ along the conservation gradient) on the Shannon diversity (A) and relative abundance (B) of saprotrophic, plant pathogenic and arbuscular mycorrhizal fungi (AMF) in bulk soil (*n* = 73). Each point represents the predicted marginal effect with the horizontal line showing 95% confidence interval from a linear mixed effect model.

**Table 1:** Summary of linear mixed-effect models testing how plant species richness and the community root trait gradients affect Shannon diversity and relative abundance of saprotrophic, plant pathogenic and arbuscular mycorrhizal fungi (see Fig. 3).

| Response | Guild | Predictor | Standardized Estimate | SE | P |
| --- | --- | --- | --- | --- | --- |
| Shannon diversity | Saprotrophs | Plant species richness (log) | 0.273 | 0.102 | **0.009** |
|  |  | Collaboration gradient ('outsourcing') | -0.010 | 0.095 | 0.920 |
|  |  | Conservation gradient ('fast') | -0.274 | 0.098 | **0.007** |
|  | Plant pathogens | Plant species richness (log) | 0.100 | 0.120 | 0.409 |
|  |  | Collaboration gradient ('outsourcing') | -0.273 | 0.113 | **0.019** |
|  |  | Conservation gradient ('fast') | 0.297 | 0.117 | **0.013** |
|  | AMF | Plant species richness (log) | 0.106 | 0.115 | 0.359 |
|  |  | Collaboration gradient ('outsourcing') | 0.302 | 0.107 | **0.006** |
|  |  | Conservation gradient ('fast') | -0.093 | 0.110 | 0.403 |
| Relative abundance | Saprotrophs | Plant species richness (log) | 0.062 | 0.111 | 0.579 |
|  |  | Collaboration gradient ('outsourcing') | 0.250 | 0.105 | **0.020** |
|  |  | Conservation gradient ('fast') | 0.166 | 0.109 | 0.132 |
|  | Plant pathogens | Plant species richness (log) | 0.190 | 0.127 | 0.138 |
|  |  | Collaboration gradient ('outsourcing') | -0.291 | 0.117 | **0.015** |
|  |  | Conservation gradient ('fast') | 0.171 | 0.121 | 0.162 |
|  | AMF | Plant species richness (log) | -0.089 | 0.127 | 0.487 |
|  |  | Collaboration gradient ('outsourcing') | 0.118 | 0.120 | 0.326 |
|  |  | Conservation gradient ('fast') | -0.001 | 0.124 | 0.995 |

The scores of each plant community (n = 73) along the first and second rotated axis in the root trait PCA were extracted and used as fixed effect in the model. Standardized estimates were obtained by z-transformation of variables prior to fitting the model. Experimental block was included as a random effect to account for spatial effects in the field site. AMF, arbuscular mycorrhizal fungi; SE, standard error.

Plant pathogenic fungi did not change in their Shannon diversity along the plant species richness gradient (Table 1, Fig. 3), suggesting that higher morphological and chemical diversity of roots at higher species richness does not increase pathogen diversity. Instead, the lower resource quality and lower host density for specialist pathogens in diverse plant communities^53,54^ might limit pathogen diversity. The result, however, is in contrast to studies on aboveground pathogens that found plant species richness to also increase pathogen richness^55^, indicating that pathogen dynamics belowground do not follow the same trends as aboveground. The root trait gradients, on the other hand, were strong predictors of pathogen diversity, with ‘outsourcing’ traits along the collaboration axis and ‘slow’ traits along the conservation axis being linked with lower pathogen diversity (Table 1, Fig. 3). The relative abundance of fungal pathogens also decreased with ‘outsourcing’ traits (Table 1, Fig. 3). These results are in line with our predictions and with studies that found traits of the collaboration axis at the species-level to be closely related to the fungal pathogen community^30,31,56^. To the best of our knowledge, our results show for the first time that these effects scale from the plant species- to the community-level, as well as from the root or rhizosphere to bulk soil. Similar to the species-level, we expect the suppression of plant pathogens by mycorrhizal symbionts to be the most likely explanation for the change along the collaboration axis^17^. Additionally, higher root diameter itself might also be a beneficial strategy against plant pathogens, as it decreases the relative root surface^57^ and therefore the potential contact points with pathogens. The decreased plant investments into defense in more resource-acquisitive plant communities along the conservation gradient^32^ likely allows more plant pathogenic fungi to colonize the plant and thus be more diverse and abundant in the soil as well.

AMF diversity was significantly linked with the collaboration axis, with higher AMF diversity found in plant communities with ‘outsourcing’ root strategies (Table 1, Fig. 3). Based on the reliance of AMF on high cortex volume and root diameter^58^, higher intra-radical mycorrhizal colonization rate and higher extra-radical hyphal length^59^ and therefore potentially also higher abundance and diversity is expected with ‘outsourcing’ roots. Generally, AMF communities in bulk soil are more diverse than in the root, as the plant only recruits a fraction of species from the available species pool in the soil^60,61^. While we did not measure mycorrhizal colonization in this study, the positive correlation with root diameter has been previously shown for a subset of species in our field site^62^ and our data now highlight that these trait-fungal relationships at the plant species-level also transfer to AMF diversity in the soil at the plant community-level. Plant species richness and the conservation axis were not related to AMF diversity (Table 1, Fig. 3). The general direction of effects on the relative abundance of AMF was similar to effects on AMF diversity, but there was no significant relationship with the collaboration axis. While we calculated AMF diversity from sequence data of the AMF-specific primers, relative abundance compared to other fungal guilds was calculated from the ITS2 sequence data, in which AMF only account for a very small portion due to the primer bias^63^. We therefore attribute these weaker effects on the relative abundance of AMF to the sequencing methods rather than ecological effects.

Overall, we found strong effects of the plant community root trait gradients on the diversity and relative abundance of fungal guilds, with each being significantly correlated with at least one trait axis. Plant species richness, however, was considerably less important than the trait axes. Specifically, we found no change in the relative abundance of any of the three fungal guilds in response to the plant species richness gradient (Table 1, Fig. 3), suggesting that the fungal guild composition of the fungal community is less sensitive to the diversity of the root system and quantity and quality of plant litter input determined by plant species richness compared to root trait axes.

### Drivers of fungal and microbial biomass

Sequencing studies have substantially advanced our knowledge of the community composition of soil microbial communities and are an indispensable part of soil ecology. Yet, the increased use of compositional sequence data has partly shifted focus away from more quantitative measures of soil microbial communities. To gain a more holistic view of the effects of plant species richness and root traits on soil fungal communities, we also quantified lipid biomarkers from soil samples.

The biomass of fine roots, a critical carbon source for the majority of soil fungi^38^, increased with plant species richness but was also significantly higher in plant communities with ‘outsourcing’ and ‘slow’ root trait strategies (Table 2, Fig. 4). Traits associated with these strategies (i.e. high root diameter and high root tissue density) generally show a positive relationship with root life span^64,65^ and can therefore enhance root standing biomass. The fungal phospholipid fatty acid (PLFA) marker 18:2ω6,9, indicative of overall fungal biomass, increased strongly with plant species richness but was not associated with the collaboration and conservation axis of root traits (Table 2, Fig. 4). The soil microbial biomass carbon, calculated from substrate-induced soil respiration, showed a similar positive effect of plant species richness but no effect of the root trait gradients (Table 2, Fig. 4). The AMF-specific neutral lipid fatty acids (NLFA) marker 16:1ω5, however, was not significantly related to either plant species richness or trait axes (Table 2, Fig. 4). Given that plant species richness was previously shown to increase carbon transport to AMF^66^ and that the root length colonized by AMF is correlated with AMF biomass in the soil^67^, it is surprising that the AMF biomarker did not show a positive relationship with ‘outsourcing’ root strategies and increasing plant species richness. NLFA, unlike PLFA, are mainly storage lipids or found in spores^68^ and are therefore not as directly relatable to fungal biomass, potentially explaining the weak effect. While our study was not designed to specifically test this, our results do not support the hypothesis that the diversity, relative abundance, or biomass of AMF mediate positive biodiversity-ecosystem functioning (BEF) relationships^69,70^.

**Fig. 4:**
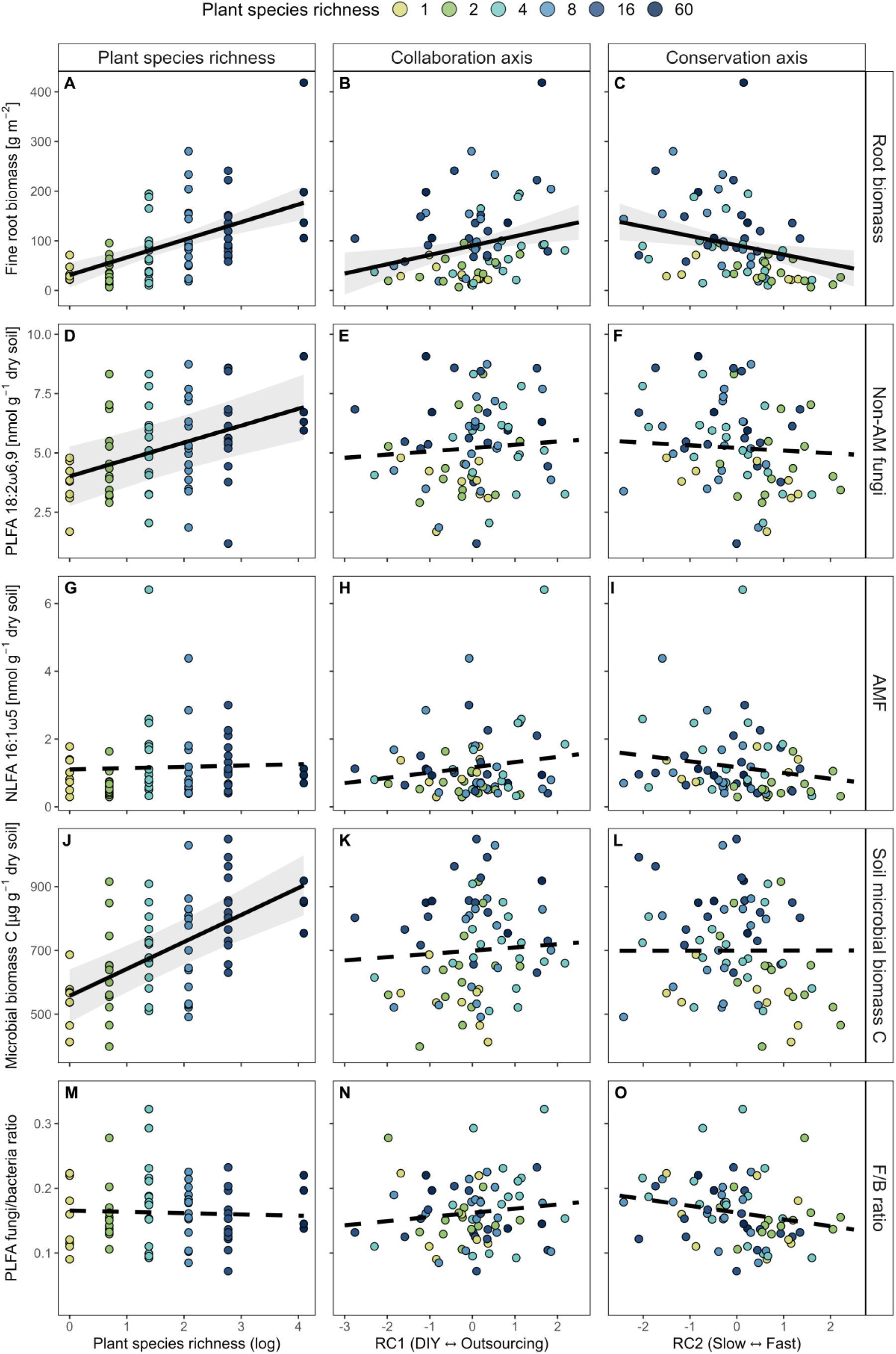
The relationship of plant species richness and root trait gradients with root biomass and soil microbial properties. Changes in root biomass, non-arbuscular mycorrhizal (non-AM) and arbuscular mycorrhizal fungi (AMF) biomarker concentration, soil microbial biomass carbon and fungal to bacterial (F/B) ratio along the plant species richness gradient and the collaboration and conservation axis of the root economics space. Each point represents one experimental plot (*n* = 70). Regression lines are based on linear mixed-effect model predictions, solid lines indicate significant relationships (P<0.05), dashed lines indicate non-significant relationships (P>0.05). The grey bands around significant regression lines depict the 95% confidence interval.

**Table 2:** Summary of results of linear mixed-effect models testing how plant species richness and the community root trait gradients affect root biomass, PLFA and NLFA biomarkers, soil microbial biomass carbon and the fungal : bacterial ratio (see Fig. 4).

| Response | Predictor | Estimate | SE | P |
| --- | --- | --- | --- | --- |
| Fine root biomass | Plant species richness (log) | 35.552 | 6.594 | **<0.001** |
|  | Collaboration gradient ('outsourcing') | 18.765 | 6.794 | **0.007** |
|  | Conservation gradient ('fast') | -18.804 | 7.011 | **0.009** |
| Non-AM fungi PLFA<br>(18:2ω6,9) | Plant species richness (log) | 0.712 | 0.164 | **<0.001** |
|  | Collaboration gradient ('outsourcing') | 0.137 | 0.174 | 0.435 |
|  | Conservation gradient ('fast') | -0.111 | 0.181 | 0.541 |
| AMF NLFA<br>(16:1ω5) | Plant species richness (log) | 0.038 | 0.116 | 0.744 |
|  | Collaboration gradient ('outsourcing') | 0.155 | 0.120 | 0.201 |
|  | Conservation gradient ('fast') | -0.171 | 0.124 | 0.173 |
| Soil microbial biomass<br>carbon | Plant species richness (log) | 84.701 | 13.316 | **<0.001** |
|  | Collaboration gradient ('outsourcing') | 10.163 | 14.115 | 0.474 |
|  | Conservation gradient ('fast') | 0.187 | 14.618 | 0.990 |
| Fungal : bacterial ratio | Plant species richness (log) | -0.002 | 0.005 | 0.710 |
|  | Collaboration gradient ('outsourcing') | 0.006 | 0.006 | 0.253 |
|  | Conservation gradient ('fast') | -0.010 | 0.006 | 0.075 |

The scores of each plant community (n = 70) along the first and second rotated axis in the root trait PCA were extracted and used as fixed effect in the model. Experimental block was included as a random effect to account for spatial effects in the field site. SE, standard error; non-AM, non-arbuscular mycorrhizal; AMF, arbuscular mycorrhizal fungi; PLFA, Phospholipid fatty acids; NLFA, neutral lipid fatty acid.

The ratio between fungal and bacterial biomass (F/B) is considered a proxy for nutrient cycling rates in soils as a higher fungal proportion decreases nutrient cycling rates and increases nutrient retention compared to bacterial-dominated communities^71,72^. We found no change in the F/B ratio along the plant species richness gradient and the collaboration axis but a marginally significant decrease with ‘fast’ root traits along the conservation axis (Table 2, Fig. 4). This aligns with previous concepts and results suggesting that bacteria benefit from the higher litter quality of ‘fast’ above- and belowground traits^72^.

Generally, the quantification of absolute abundances or biomass of individual fungal guilds is not possible in the same way as for relative abundances. Approaches using qPCR methods can quantify gene copies under certain circumstances but are sensitive to biases during the DNA extraction or require standardization^73,74^. Since the PLFA biomarker 18:2ω6,9 is largely determined by the biomass of Ascomycota and Basidiomycota^75^, we used it as an indicator of fungal biomass of non-arbuscular mycorrhizal fungi, which include saprotrophic as well as pathogenic fungi. Because PLFA biomarkers and sequencing rely on very different components of the fungal community, the results are not directly comparable. However, as the relative abundance of saprotrophs and plant pathogens changes along the root trait axes but the fungal biomass is not affected, we conclude that the trait axes affect the ratio of guilds in the fungal community, but not the biomass of individual guilds. Studies using a combined qualitative (e.g. sequencing) and quantitative (e.g. PLFA and respiration) approach provide valuable opportunities to overcome the limitations of the compositional nature of sequencing data in the absence of appropriate qPCR methods.

### Conclusions

In summary, our study demonstrates that in experimental grassland communities, fine root functional strategies of the root economics space scale from the species-level to the community-level. We further demonstrate that these functional strategies of plant communities structure the guild composition of soil fungal communities, with saprotrophic, plant pathogenic and arbuscular mycorrhizal varying in diversity or relative abundance depending on the root traits of the plant community. Plant species richness, however, is only a weak driver of the fungal and microbial community composition but drives microbial and fungal biomass in the soil. Ultimately, root trait gradients drive the soil fungal guild composition, but plant species richness controls the fungal biomass. These contrasting results on the role of plant species richness and root trait gradients highlight that a diversity of mechanisms need to be considered in future predictions of how changes in plant communities will affect soil biodiversity and functioning.

## Methods

### Sampling design and soil collection

We conducted our study at the Jena Experiment, a large-scale long-term biodiversity experiment. The area is located on a former arable field near the river Saale 51° N, 11° E, 135 m above sea level^76^. In 2002, experimental plots varying in sown plant species richness from 1 to 2, 4, 8, 16, and 60 species were set up. The plots are grouped in four blocks to account for the differences in initial soil conditions^76^. The initial size of the plots was 10 × 10 m before being reduced to 6 × 5.5 m in 2010. The plots are mown twice a year with the plant material being removed and not fertilized. They are further weeded manually two to three times per year to maintain the designed plant communities. There was no resowing of any plant species, leading to local extinction of some species. However, sown and realized plant species richness are strongly positively correlated^77^.

For root trait measurements and DNA extraction, we took 4 soil cores (3.5 cm diameter, 5 cm depth) between May 31^st^ and June 11^th^ 2021 in four locations across each plot. The soil cores were stored at 4 °C until final preparation (no longer than 24 h after sampling). The four soil samples were pooled, and a subsample was collected for the DNA extraction and frozen at –20 °C immediately. For respiration measurement and fatty acid analysis, we took four soil cores (2 cm diameter, 10 cm depth) in each plot between June 14^th^ and 24^th^ 2021, and stored them at 4 °C until measurements were done.

### Root trait measurements

Of the 80 plots in the Jena Experiment, 7 plots did not contain any of the sown plant species or did not yield enough root material to measure root traits and were therefore not further considered in our sampling. The pooled soil cores for trait measurements were soaked in water for around 15 minutes and then washed with tap water over a sieve and manually cleaned. Coarse roots with a diameter larger than 2 mm were manually removed from the sample. A random subset of fine roots was scanned and measured using an Epson Expression 11000XL (Epson, Tokyo, Japan) flatbed scanner at 600 dpi and the software RhizoVision Explorer^78^ to quantify root length, root diameter, and root volume. The scanned roots were weighed, dried (48 h at 70 °C), and then weighed again for dry mass. Specific root length (SRL) was calculated as root length : dry mass and root tissue density (RTD) as root dry mass : root volume. Fine root biomass in g/m^2^ was calculated based on root dry mass per area of the soil cores. The roots were freeze-dried and ground using a zirconium kit in a ball mill (MM400, Retsch, Haan, Germany). For 46 random samples, relative nitrogen content (RN, % of dry weight) was quantified using an elemental analyzer (Elementar vario ELII, Hanau, Germany) at the Max-Planck-Institute for Biogeochemistry in Jena, Germany. All samples were freeze-dried again to measure near-infrared spectra (NIR) in the range of 9090–4000cm^−1^ at 8cm^−1^ resolution in transmission mode (Multi-Purpose FT-NIR-Analyzer, Bruker Corporation, Billerica, USA). For each sample, five independent measurements were averaged. We converted transmission to absorbance as log_10_(1/Transmission) and used it in combination with a bootstrapped CARS-PLSR procedure^79^ to predict nitrogen content for the remaining 26 samples.

The sampling of root traits at the community-level, as used in our study, differs from the more common species-level sampling. In grasslands, our method allows for faster measurement of community traits, while still including plasticity in species traits that would not be captured if a species was only sampled in some communities. Additionally, belowground community weighted-mean traits are usually calculated based on aboveground plant community composition, assuming similar belowground composition. Disproportional above- or belowground allocation at the species-level can therefore cause a misrepresentation of community weighted-mean traits, whereas they do not affect our directly measured community traits. At the same time, every plant community is only described by one value per trait, rather than a weighted mean calculated from individual species’ traits, and it is therefore not possible to quantify functional diversity.

### Fungal amplicon sequencing

We extracted genomic DNA from 0.25 - 0.3 g of thawed and homogenized soil using the Quick-DNA Fecal/Soil Microbe Miniprep Kit (Zymo Research Europe, Freiburg, Germany) following the manufacturer’s instructions. We measured the DNA content using a NanoDrop 2000c spectrophotometer (Thermo Fisher Scientific, Dreieich, Germany) and stored the DNA at −20 °C until amplification. We amplified the internal transcribed spacer (ITS) region 2 using ITS4 and ITS7:ITS7o primers^80,81^. Library preparation and sequencing followed the protocol described in Hennecke *et al.*^17^. In short, three positive PCR products were pooled, purified with AMPure XP beads (Beckman Coulter), and indexed with the Nextera XT Illumina Index Kit (Illumina Inc., San Diego, USA). The samples were then pooled to equal molarity and the paired-end sequencing of 2 × 300 bp was performed using a MiSeq Reagent Kit v3 on an Illumina MiSeq System at the Department of Soil Ecology of the Helmholtz-Centre for Environmental Research (UFZ, Halle/Saale, Germany). To overcome the amplification bias against AMF with ITS primers^63^, we additionally sequenced AMF. For this, the SSU of AMF was amplified using a nested PCR with primer pair Glomer 1536 / WT0 for the first PCR and NS31 / AML2 for the second PCR according to Wahdan *et al.*^82^ and the sequencing was performed as for the ITS sequencing. Detailed description of the sequencing of AMF communities is described in Albracht *et al.*^83^.

The processing of sequences was executed using the snakemake implementation *dadasnake*^84^ of the *DADA2* pipeline^85^. To summarize, *cutadapt*^86^ was used to trim the primer sequences from raw reads. Quality trimming was performed with a minimum length of 70 bp for ITS and 260 bp (fwd) / 210 bp (rvs) for AMF, truncation of reads at positions with a PHRED score below 15 for ITS, and exclusion of reads with an expected error higher than 2. The identification of exact sequence variants (Amplicon Sequence Variants, ASVs) included merging read pairs with a minimum overlap of 15 bp (ITS) and 12 bp (AMF) and a maximum of three (ITS) and zero (AMF) mismatches. Chimeras were filtered using DADA2’s ’consensus’ algorithm. Taxonomic classification was conducted using the *mothur*^87^ implementation of the Bayesian classifier against the UNITE v8.2 database^88^ (ITS) and SILVA v138 SSUref database^89^ for AMF. Non-fungal ASVs (for ITS sequences) and non-Glomeromycotinian ASVs (for AMF sequences) were discarded. For ITS data, ASVs were then assigned to putative fungal guilds based on their taxonomic annotation and the FungalTraits database^20^. Further, all Glomeromycotinian ASVs in the ITS data were assigned to be arbuscular mycorrhizal. For the AMF data, ASVs were blasted against the MaarjAM database^90^ and assigned to virtual taxa (VTX). ASVs without VTX assignment were extracted, singletons removed, and used for the construction of a maximum likelihood phylogenetic tree based on a general time-reversible, discrete gamma (GTR+G) model using raxML^91^ and FasttreeMP^92^. Consequently, these ASVs were associated with custom virtual taxa (VTC) characterized by cophenetic distances below 0.03. For the analysis of fungal diversity, the dataset was rarefied to address any potential impact of sequencing depth on ASV richness (Supplementary Fig. S1).

### Lipid fatty acid and respiration measurement

Soil fungal biomass was determined using phospholipid fatty acids (PLFA) analysis. We extracted PLFAs from 5 g of soil following Frostegård *et al.*^93^ and fractioned them into PLFAs, neutral lipid fatty acids (NLFA), and glycolipids. PLFAs and NLFA were then measured using a gas-chromatograph (GC-FID Clarus 500; PerkinElmer Corporation, Norwalk, USA) with an Elite-5 column (PerkinElmer Corporation, Norwalk, USA). PLFA and NLFA concentrations were calculated from the internal standard C19:0 (Methylnonadecanoat). Based on the classification of Ruess and Chamberlain^94^, we used 18:2ω6,9 PLFA marker as a measure of fungal biomass and the 16:1ω5 NLFA marker as a measure of AMF biomass. For the fungal : bacteria (F/B) ratio, the sum of both fungal markers was divided by the sum of the bacterial PLFA markers a15:0, i15:0, i16:0, i17:0, 16:1ω7, cy17:0, and cy19:0. For three samples, chromatograms showed very high peaks suggesting erratic measurements but could not be repeated due to small sample amount and were therefore excluded from the analysis. As recent studies raised the issue that plant biomass can also contribute to 18:2ω6,9 PLFA^95^, we further quantified soil microbial biomass carbon (C_mic_) using an O2-micro-compensation apparatus^96^. The soil was sieved at 2 mm to remove stones, large organic materials, and larger organisms, and added watery glucose solution to determine the maximal initial respiratory response (MIRR). C_mic_ was calculated from MIRR following Beck *et al.*^97^. Soil water content was estimated via drying after the end of measurements.

### Statistical analysis

Data analyses were conducted in R v.4.3.2^98^. We used a principal component analysis (PCA) of the root traits RD, SRL, RN, and RTD, followed by varimax rotation for better interpretability of the components and inversing the scores and loadings to match the direction of trait gradients in Bergmann *et al.*^11^. Rotated components (RC) 1 and 2 of the PCA captured the root economics space’s main axes (Fig. 1), representing the collaboration and conservation axes. Subsequent analyses used the scores from these axes to examine the effects on fungal communities. The unrotated PCA is shown in Supplementary Fig. S2.

Fungal abundance and taxonomy were organized in a phyloseq object^99^. Read numbers of ASVs with a primary lifestyle as litter saprotrophs, soil saprotrophs, wood saprotrophs, and unspecified saprotrophs were summed for the total number of reads of saprotrophs. Shannon diversity was calculated within the three fungal guilds (saprotrophs, plant pathogens, and AMF). Relative guild abundance was calculated as the number of reads per guild relative to the total number of reads. A broad description of the sequenced fungal ITS2 community is presented in Supplementary Methods S1.

To test how plant species richness affected the root trait axes, we used two separate linear mixed models with RC1 and RC2 as the response and plant species richness (log) as a fixed effect and the experimental block as a random term. We tested how plant species richness and root traits axes are linked with fungal guild diversity and relative abundance, as well as root biomass, PLFA and NLFA biomarker concentration, F/B ratio, and soil microbial carbon, using a linear mixed-effect models of the lme4 package^100^ and tested for significance using lmerTest^101^ with type Ⅲ sum of squares. Log-scaled sown plant species richness and RC1 and RC2 were included as the fixed effects and the experimental block as a random term to account for any spatial effects of the field site. Standardized effect sizes were extracted using z-transformed model variables. In case the random terms did not explain any variance, linear regression was used instead. Collinearity was checked for all models but was considered unproblematic with variance inflation factors well below 2 for all models. An additional analysis with sequential fitting of variables is presented in Supplementary Table S2.

## Acknowledgements

We thank Anne Ebeling and Anna-Maria Madaj for coordination of the Jena Experiment, as well as the technical staff and numerous student helpers for the maintenance and measurements. We are grateful to François Buscot for granting access to the sequencing facilities at the Helmholtz-Centre for Environmental Research (Halle/Saale), and to Beatrix Schnabel for her technical expertise in sample preparation and sequencing. Data processing was performed at the High-Performance Computing (HPC) Cluster EVE, a joint effort of the Helmholtz Centre for Environmental Research–UFZ and the German Centre for Integrative Biodiversity Research (iDiv) Halle-Jena-Leipzig, and we thank Christian Krause and the other administrators for excellent support. We thank Stefan Scheu for the measurement of PLFA/NLFA biomarkers and Simone Cesarz for helping with the analysis of the PLFA/NLFA data. Markus Lange gratefully acknowledges funding by the Zwillenberg-Tietz Foundation. We acknowledge the support of iDiv funded by the German Research Foundation (DFG–FZT 118, 202548816). The Jena Experiment is funded by the DFG (FOR 5000).

## Author contributions

J.H., J.B., N.E., A.H.-B., T.W.K., M.L., L.M. and A.W. conceived the idea of the study. J.H., L.B., C.A., A.A., L.H., Y.P., A.R. and M.D.S. collected the data; J.H. and A.H.-B. processed the sequence data and J.H. analyzed the data. J.H. led the writing of the manuscript and all authors contributed to reviewing and editing.

## Data availability

The data associated with this study will be published with a unique accession number. The raw Illumina sequences generated in this study are available in the NCBI Sequence Read Archive (BioProject: PRJNA1074103 and PRJNA988299).

## Code availability

The code used for the analyses and to produce the figures will be deposited together with the data.

## References

1. Tedersoo, L. et al. Global diversity and geography of soil fungi. Science 346, 1256688 (2014).

2. Tedersoo, L., Bahram, M. & Zobel, M. How mycorrhizal associations drive plant population and community biology. Science 367, eaba1223 (2020).

3. Wagg, C., Bender, S. F., Widmer, F. & van der Heijden, M. G. A. Soil biodiversity and soil community composition determine ecosystem multifunctionality. Proc. Natl. Acad. Sci. 111, 5266–5270 (2014).

4. Schmidt, R., Mitchell, J. & Scow, K. Cover cropping and no-till increase diversity and symbiotroph:saprotroph ratios of soil fungal communities. Soil Biol. Biochem. 129, 99–109 (2019).

5. Lutz, S. et al. Soil microbiome indicators can predict crop growth response to large-scale inoculation with arbuscular mycorrhizal fungi. Nat. Microbiol. 8, 2277–2289 (2023).

6. Jansson, J. K., McClure, R. & Egbert, R. G. Soil microbiome engineering for sustainability in a changing environment. Nat. Biotechnol. 41, 1716–1728 (2023).

7. Scherber, C. et al. Bottom-up effects of plant diversity on multitrophic interactions in a biodiversity experiment. Nature 468, 553–556 (2010).

8. Zak, D. R., Holmes, W. E., White, D. C., Peacock, A. D. & Tilman, D. Plant Diversity, Soil Microbial Communities, and Ecosystem Function: Are There Any Links? Ecology 84, 2042–2050 (2003).

9. Reynolds, H. L., Packer, A., Bever, J. D. & Clay, K. Grassroots Ecology: Plant–Microbe– Soil Interactions as Drivers of Plant Community Structure and Dynamics. Ecology 84, 2281–2291 (2003).

10. Bever, J. D. et al. Rooting theories of plant community ecology in microbial interactions. Trends Ecol. Evol. 25, 468–478 (2010).

11. Bergmann, J. et al. The fungal collaboration gradient dominates the root economics space in plants. Sci. Adv. 6, eaba3756 (2020).

12. Wardle, D. A. et al. Ecological Linkages Between Aboveground and Belowground Biota. Science 304, 1629–1633 (2004).

13. Bergmann, J., Ryo, M., Prati, D., Hempel, S. & Rillig, M. C. Root traits are more than analogues of leaf traits: the case for diaspore mass. New Phytol. 216, 1130–1139 (2017).

14. Kong, D. et al. Leading dimensions in absorptive root trait variation across 96 subtropical forest species. New Phytol. 203, 863–872 (2014).

15. Han, M. & Zhu, B. Linking root respiration to chemistry and morphology across species. Glob. Change Biol. 27, 190–201 (2021).

16. Spitzer, C. M. et al. Root trait–microbial relationships across tundra plant species. New Phytol. 229, 1508–1520 (2021).

17. Hennecke, J. et al. Responses of rhizosphere fungi to the root economics space in grassland monocultures of different age. New Phytol. 240, 2035–2049 (2023).

18. Lachaise, T. et al. Soil conditions drive below-ground trait space in temperate agricultural grasslands. J. Ecol. 110, 1189–1200 (2022).

19. Da, R., Fan, C., Zhang, C., Zhao, X. & von Gadow, K. Are absorptive root traits good predictors of ecosystem functioning? A test in a natural temperate forest. New Phytol. 239, 75–86 (2023).

20. Põlme, S. et al. FungalTraits: a user-friendly traits database of fungi and fungus-like stramenopiles. Fungal Divers. 105, 1–16 (2021).

21. Waldrop, M. P., Zak, D. R., Blackwood, C. B., Curtis, C. D. & Tilman, D. Resource availability controls fungal diversity across a plant diversity gradient. Ecol. Lett. 9, 1127– 1135 (2006).

22. Albornoz, F. E., Prober, S. M., Ryan, M. H. & Standish, R. J. Ecological interactions among microbial functional guilds in the plant-soil system and implications for ecosystem function. Plant Soil 476, 301–313 (2022).

23. Graham, E. B. et al. Microbes as Engines of Ecosystem Function: When Does Community Structure Enhance Predictions of Ecosystem Processes? Front. Microbiol. 7, (2016).

24. van der Heijden, M. G. A., Bardgett, R. D. & van Straalen, N. M. The unseen majority: soil microbes as drivers of plant diversity and productivity in terrestrial ecosystems. Ecol. Lett. 11, 296–310 (2008).

25. Ferlian, O., Wirth, C. & Eisenhauer, N. Leaf and root C-to-N ratios are poor predictors of soil microbial biomass C and respiration across 32 tree species. Pedobiologia 65, 16–23 (2017).

26. Barberán, A. et al. Relating belowground microbial composition to the taxonomic, phylogenetic, and functional trait distributions of trees in a tropical forest. Ecol. Lett. 18, 1397–1405 (2015).

27. de Vries, F. T. et al. Abiotic drivers and plant traits explain landscape-scale patterns in soil microbial communities. Ecol. Lett. 15, 1230–1239 (2012).

28. Weigelt, A. et al. An integrated framework of plant form and function: the belowground perspective. New Phytol. 232, 42–59 (2021).

29. Delavaux, C. S., Smith-Ramesh, L. M. & Kuebbing, S. E. Beyond nutrients: a meta-analysis of the diverse effects of arbuscular mycorrhizal fungi on plants and soils. Ecology 98, 2111– 2119 (2017).

30. Semchenko, M. et al. Fungal diversity regulates plant-soil feedbacks in temperate grassland. Sci. Adv. 4, eaau4578 (2018).

31. Wang, Y., Wang, J., Qu, M. & Li, J. Root attributes dominate the community assembly of soil fungal functional guilds across arid inland river basin. Front. Microbiol. 13, (2022).

32. Coley, P. D., Bryant, J. P. & Chapin, F. S. Resource Availability and Plant Antiherbivore Defense. Science 230, 895–899 (1985).

33. Otsing, E., Barantal, S., Anslan, S., Koricheva, J. & Tedersoo, L. Litter species richness and composition effects on fungal richness and community structure in decomposing foliar and root litter. Soil Biol. Biochem. 125, 328–339 (2018).

34. Francioli, D. et al. Plant functional group drives the community structure of saprophytic fungi in a grassland biodiversity experiment. Plant Soil 461, 91–105 (2021).

35. Chen, Y.-L. et al. Plant diversity represents the prevalent determinant of soil fungal community structure across temperate grasslands in northern China. Soil Biol. Biochem. 110, 12–21 (2017).

36. Hooper, D. U. et al. Interactions between Aboveground and Belowground Biodiversity in Terrestrial Ecosystems: Patterns, Mechanisms, and Feedbacks. BioScience 50, 1049–1061 (2000).

37. Steinauer, K., Chatzinotas, A. & Eisenhauer, N. Root exudate cocktails: the link between plant diversity and soil microorganisms? Ecol. Evol. 6, 7387–7396 (2016).

38. Eisenhauer, N. et al. Root biomass and exudates link plant diversity with soil bacterial and fungal biomass. Sci. Rep. 7, 44641 (2017).

39. Tilman, D. et al. Diversity and Productivity in a Long-Term Grassland Experiment. Science 294, 843–845 (2001).

40. van der Heijden, M. G. A. et al. Mycorrhizal fungal diversity determines plant biodiversity, ecosystem variability and productivity. Nature 396, 69–72 (1998).

41. Yang, T. et al. Soil fungal diversity in natural grasslands of the Tibetan Plateau: associations with plant diversity and productivity. New Phytol. 215, 756–765 (2017).

42. Dassen, S. et al. Differential responses of soil bacteria, fungi, archaea and protists to plant species richness and plant functional group identity. Mol. Ecol. 26, 4085–4098 (2017).

43. Chen, W. et al. Fertility-related interplay between fungal guilds underlies plant richness– productivity relationships in natural grasslands. New Phytol. 226, 1129–1143 (2020).

44. Antoninka, A., Reich, P. B. & Johnson, N. C. Seven years of carbon dioxide enrichment, nitrogen fertilization and plant diversity influence arbuscular mycorrhizal fungi in a grassland ecosystem. New Phytol. 192, 200–214 (2011).

45. Eisenhauer, N. et al. Plant diversity effects on soil microorganisms support the singular hypothesis. Ecology 91, 485–496 (2010).

46. Keesing, F., Holt, R. D. & Ostfeld, R. S. Effects of species diversity on disease risk. Ecol. Lett. 9, 485–498 (2006).

47. Mulder, C., Jumpponen, A., Högberg, P. & Huss-Danell, K. How plant diversity and legumes affect nitrogen dynamics in experimental grassland communities. Oecologia 133, 412–421 (2002).

48. van Ruijven, J. & Berendse, F. Diversity–productivity relationships: Initial effects, long-term patterns, and underlying mechanisms. Proc. Natl. Acad. Sci. 102, 695–700 (2005).

49. Mommer, L. et al. Diversity effects on root length production and loss in an experimental grassland community. Funct. Ecol. 29, 1560–1568 (2015).

50. Ravenek, J. M. et al. Long-term study of root biomass in a biodiversity experiment reveals shifts in diversity effects over time. Oikos 123, 1528–1536 (2014).

51. Chen, H. et al. Plant species richness negatively affects root decomposition in grasslands. J. Ecol. 105, 209–218 (2017).

52. Hunt, H. W. et al. The detrital food web in a shortgrass prairie. Biol. Fertil. Soils 3, 57–68 (1987).

53. Ampt, E. A. et al. Deciphering the interactions between plant species and their main fungal root pathogens in mixed grassland communities. J. Ecol. 110, 3039–3052 (2022).

54. Wang, G. et al. Dilution of specialist pathogens drives productivity benefits from diversity in plant mixtures. Nat. Commun. 14, 8417 (2023).

55. Rottstock, T., Joshi, J., Kummer, V. & Fischer, M. Higher plant diversity promotes higher diversity of fungal pathogens, while it decreases pathogen infection per plant. Ecology 95, 1907–1917 (2014).

56. Sweeney, C. J., de Vries, F. T., Dongen, B. E. & Bardgett, R. D. Root traits explain rhizosphere fungal community composition among temperate grassland plant species. New Phytol. 229, 1492–1507 (2021).

57. McCormack, M. L. & Iversen, C. M. Physical and Functional Constraints on Viable Belowground Acquisition Strategies. Front. Plant Sci. 10, (2019).

58. Brundrett, M. C. Coevolution of roots and mycorrhizas of land plants. New Phytol. 154, 275–304 (2002).

59. Gryndler, M. et al. Organic and mineral fertilization, respectively, increase and decrease the development of external mycelium of arbuscular mycorrhizal fungi in a long-term field experiment. Mycorrhiza 16, 159–166 (2006).

60. Johnson, D. et al. Plant communities affect arbuscular mycorrhizal fungal diversity and community composition in grassland microcosms. New Phytol. 161, 503–515 (2004).

61. Hempel, S., Renker, C. & Buscot, F. Differences in the species composition of arbuscular mycorrhizal fungi in spore, root and soil communities in a grassland ecosystem. Environ. Microbiol. 9, 1930–1938 (2007).

62. Bassi, L. et al. Uncovering the secrets of monoculture yield decline: trade-offs between leaf and root chemical and physical defence traits in a grassland experiment. Oikos 2024, e10061 (2024).

63. Tedersoo, L. et al. Shotgun metagenomes and multiple primer pair-barcode combinations of amplicons reveal biases in metabarcoding analyses of fungi. MycoKeys 10, 1–43 (2015).

64. Luke McCormack, M., Adams, T. S., Smithwick, E. A. H. & Eissenstat, D. M. Predicting fine root lifespan from plant functional traits in temperate trees. New Phytol. 195, 823–831 (2012).

65. Wahl, S. & Ryser, P. Root tissue structure is linked to ecological strategies of grasses. New Phytol. 148, 459–471 (2000).

66. Mellado-Vázquez, P. G. et al. Plant diversity generates enhanced soil microbial access to recently photosynthesized carbon in the rhizosphere. Soil Biol. Biochem. 94, 122–132 (2016).

67. Barceló, M. et al. The abundance of arbuscular mycorrhiza in soils is linked to the total length of roots colonized at ecosystem level. PLOS ONE 15, e0237256 (2020).

68. Gorka, S. et al. Beyond PLFA: Concurrent extraction of neutral and glycolipid fatty acids provides new insights into soil microbial communities. Soil Biol. Biochem. 187, 109205 (2023).

69. Eisenhauer, N., et al. Biotic Interactions as Mediators of Context-Dependent Biodiversity-Ecosystem Functioning Relationships. Res. Ideas Outcomes 8, e85873- (2022).

70. Eisenhauer, N., Reich, P. B. & Scheu, S. Increasing plant diversity effects on productivity with time due to delayed soil biota effects on plants. Basic Appl. Ecol. 13, 571–578 (2012).

71. Wardle, D. A. Communities and Ecosystems: Linking the Aboveground and Belowground Components. (Princeton University Press, Princeton, N.J, 2002).

72. Bardgett, R. D. Plant trait-based approaches for interrogating belowground function. Biol. Environ. Proc. R. Ir. Acad. 117B, 1–13 (2017).

73. Feinstein, L. M., Sul, W. J. & Blackwood, C. B. Assessment of Bias Associated with Incomplete Extraction of Microbial DNA from Soil. Appl. Environ. Microbiol. 75, 5428– 5433 (2009).

74. Baldrian, P. et al. Estimation of fungal biomass in forest litter and soil. Fungal Ecol. 6, 1– 11 (2013).

75. Klamer, M. & Bååth, E. Estimation of conversion factors for fungal biomass determination in compost using ergosterol and PLFA 18:2ω6,9. Soil Biol. Biochem. 36, 57–65 (2004).

76. Roscher, C. et al. The role of biodiversity for element cycling and trophic interactions: an experimental approach in a grassland community. Basic Appl. Ecol. 5, 107–121 (2004).

77. Weisser, W. W. et al. Biodiversity effects on ecosystem functioning in a 15-year grassland experiment: Patterns, mechanisms, and open questions. Basic Appl. Ecol. 23, 1–73 (2017).

78. Seethepalli, A. et al. RhizoVision Explorer: open-source software for root image analysis and measurement standardization. AoB PLANTS 13, (2021).

79. Richter, R. & Bassi, L. Bagging-CARS-PLS model R code. Jena Experiment Information System 10.25829/D6F2-ZP34 (2023).

80. Ihrmark, K. et al. New primers to amplify the fungal ITS2 region - evaluation by 454-sequencing of artificial and natural communities. FEMS Microbiol. Ecol. 82, 666–677 (2012).

81. Kohout, P. et al. Comparison of commonly used primer sets for evaluating arbuscular mycorrhizal fungal communities: Is there a universal solution? Soil Biol. Biochem. 68, 482– 493 (2014).

82. Wahdan, S. F. M. et al. Organic agricultural practice enhances arbuscular mycorrhizal symbiosis in correspondence to soil warming and altered precipitation patterns. Environ. Microbiol. 23, 6163–6176 (2021).

83. Albracht, C. et al. Common soil history is more important than plant history for arbuscular mycorrhizal community assembly in an experimental grassland diversity gradient. Preprint at 10.1101/2024.03.14.585138 (2024).

84. Weißbecker, C., Schnabel, B. & Heintz-Buschart, A. Dadasnake, a Snakemake implementation of DADA2 to process amplicon sequencing data for microbial ecology. GigaScience 9, 1–8 (2020).

85. Callahan, B. J. et al. DADA2: High-resolution sample inference from Illumina amplicon data. Nat. Methods 13, 581–583 (2016).

86. Martin, M. Cutadapt removes adapter sequences from high-throughput sequencing reads. EMBnet.journal 17, 10–12 (2011).

87. Schloss, P. D. et al. Introducing mothur: Open-source, platform-independent, community-supported software for describing and comparing microbial communities. Appl. Environ. Microbiol. 75, 7537–7541 (2009).

88. Nilsson, R. H. et al. The UNITE database for molecular identification of fungi: handling dark taxa and parallel taxonomic classifications. Nucleic Acids Res. 47, D259–D264 (2019).

89. Quast, C. et al. The SILVA ribosomal RNA gene database project: improved data processing and web-based tools. Nucleic Acids Res. 41, D590–D596 (2013).

90. Öpik, M. et al. The online database MaarjAM reveals global and ecosystemic distribution patterns in arbuscular mycorrhizal fungi (Glomeromycota). New Phytol. 188, 223–241 (2010).

91. Stamatakis, A. RAxML version 8: a tool for phylogenetic analysis and post-analysis of large phylogenies. Bioinformatics 30, 1312–1313 (2014).

92. Price, M. N., Dehal, P. S. & Arkin, A. P. FastTree 2 – Approximately Maximum-Likelihood Trees for Large Alignments. PLOS ONE 5, e9490 (2010).

93. Frostegård, Å., Tunlid, A. & Bååth, E. Microbial biomass measured as total lipid phosphate in soils of different organic content. J. Microbiol. Methods 14, 151–163 (1991).

94. Ruess, L. & Chamberlain, P. M. The fat that matters: Soil food web analysis using fatty acids and their carbon stable isotope signature. Soil Biol. Biochem. 42, 1898–1910 (2010).

95. Joergensen, R. G. Phospholipid fatty acids in soil—drawbacks and future prospects. Biol. Fertil. Soils 58, 1–6 (2022).

96. Scheu, S. Automated measurement of the respiratory response of soil microcompartments: Active microbial biomass in earthworm faeces. Soil Biol. Biochem. 24, 1113–1118 (1992).

97. Beck, T. et al. An inter-laboratory comparison of ten different ways of measuring soil microbial biomass C. Soil Biol. Biochem. 29, 1023–1032 (1997).

98. R Core Team. R: A Language and Environment for Statistical Computing. R Foundation for Statistical Computing, Vienna, Austria. URL https://www.R-project.org/. (2023).

99. McMurdie, P. J. & Holmes, S. phyloseq: An R Package for Reproducible Interactive Analysis and Graphics of Microbiome Census Data. PLOS ONE 8, e61217 (2013).

100. Bates, D., Mächler, M., Bolker, B. & Walker, S. Fitting Linear Mixed-Effects Models Using lme4. J. Stat. Softw. 67, 1–48 (2015).

101. Kuznetsova, A., Brockhoff, P. B. & Christensen, R. H. B. lmerTest Package: Tests in Linear Mixed Effects Models. J. Stat. Softw. 82, 1–26 (2017).

