## Supplementary Information for "Plant species richness and the root economics space drive soil fungal communities"

**Supplementary Table S1: Summary of ANOVA results for the effect of plant species richness on the rotated components of the root trait PCA and on the individual fine root traits.** The scores of each plant community (n = 73) along the first and second rotated component (RC) in the root trait PCA were extracted and used as response in the model. Experimental block was included as a random effect to account for spatial effects in the field site. df_error_, error degrees of freedom; SE, standard error.

|  | **Effect of plant species richness (log)** | | | | |
| --- | --- | --- | --- | --- | --- |
| **Response** | **df_error_** | **Coefficient** | **SE** | ***F*** | ***p*** |
| RC1 (Collaboration gradient) | 68.301 | 0.092 | 0.110 | 0.704 | 0.404 |
| RC2 (Conservation gradient) | 68.416 | -0.246 | 0.106 | 5.412 | **0.023** |
| Root diameter | 72.000 | 0.004 | 0.004 | 0.925 | 0.339 |
| Specific root length | 69.180 | -11.678 | 8.121 | 2.068 | 0.155 |
| Root nitrogen | 71.000 | -0.176 | 0.046 | 14.417 | **<0.001** |
| Root tissue density | 69.328 | 0.003 | 0.003 | 0.8616 | 0.357 |

**Supplementary Table S2: Summary of ANOVA with type 1 sum-of-squares results for fungal diversity and relative abundance of guilds.** The scores of each plant community (n = 73) along the first and second rotated axis in the root trait PCA were extracted and used as fixed effect in the model. Experimental block was included as a random effect to account for spatial effects in the field site. AMF, arbuscular mycorrhizal fungi; df_error_, error degrees of freedom; SE, standard error.

| **Response** | **Guild** | **Predictor** | **df** | **df_error_** | ***F*** | ***p*** |
| --- | --- | --- | --- | --- | --- | --- |
| Shannon diversity | Saprotrophs | Plant species richness (log) | 1 | 66.576 | 12.454 | **0.001** |
|  |  | Collaboration gradient (‘outsourcing’) | 1 | 69.000 | 0.028 | 0.867 |
|  |  | Conservation gradient (‘fast’) | 1 | 68.869 | 7.785 | **0.007** |
|  | Plant pathogens | Plant species richness (log) | 1 | 66.252 | 0.002 | 0.968 |
|  |  | Collaboration gradient (‘outsourcing’) | 1 | 68.613 | 5.531 | **0.022** |
|  |  | Conservation gradient (‘fast’) | 1 | 68.828 | 6.459 | **0.013** |
|  | AMF | Plant species richness (log) | 1 | 66.533 | 2.079 | 0.154 |
|  |  | Collaboration gradient (‘outsourcing’) | 1 | 67.964 | 7.861 | **0.007** |
|  |  | Conservation gradient (‘fast’) | 1 | 66.026 | 0.708 | 0.403 |
| Relative abundance | Saprotrophs | Plant species richness (log) | 1 | 66.136 | 0.162 | 0.689 |
|  |  | Collaboration gradient (‘outsourcing’) | 1 | 67.629 | 5.832 | **0.018** |
|  |  | Conservation gradient (‘fast’) | 1 | 67.830 | 2.323 | 0.132 |
|  | Plant pathogens | Plant species richness (log) | 1 | 69.000 | 0.917 | 0.342 |
|  |  | Collaboration gradient (‘outsourcing’) | 1 | 69.000 | 5.996 | **0.017** |
|  |  | Conservation gradient (‘fast’) | 1 | 69.000 | 2.001 | 0.162 |
|  | AMF | Plant species richness (log) | 1 | 66.209 | 0.402 | 0.528 |
|  |  | Collaboration gradient (‘outsourcing’) | 1 | 68.333 | 0.981 | 0.326 |
|  |  | Conservation gradient (‘fast’) | 1 | 68.572 | 0.000 | 0.995 |

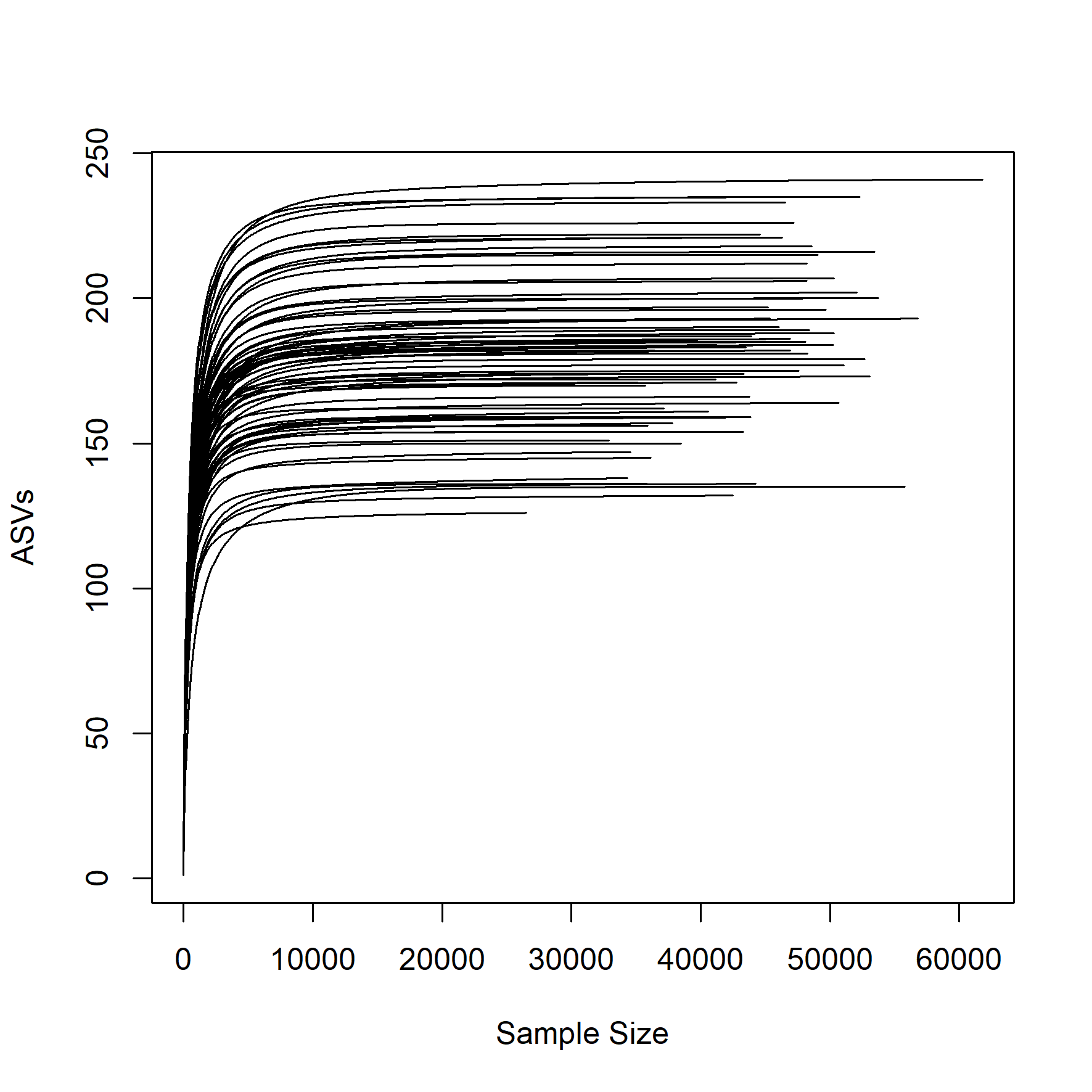

**Supplementary Fig. S1: Rarefaction curves of the ITS2 sequence data.** Each line represents one of the 73 samples, showing increasing number of amplicon sequence variants (ASVs) with the number of reads until saturation is reached.

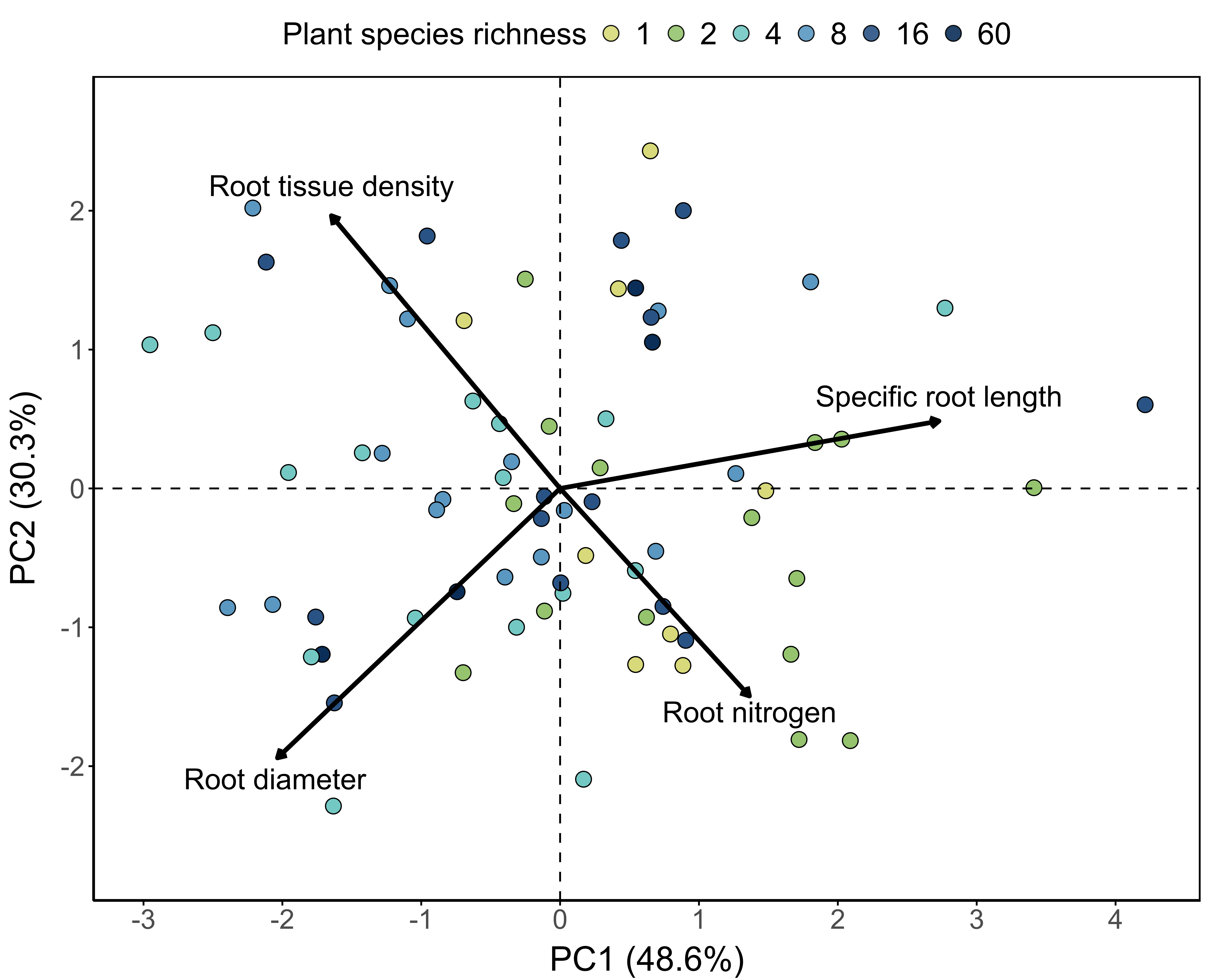
 **Supplementary Fig. S2: Unrotated PCA of the community root traits.** Each point represents a plant community (*n* = 73). Points are color-coded for plant species richness of the plot. RC, rotated component.

**Supplementary Methods S1: Description of the final ITS2 dataset.**

Our filtered ITS2 sequence dataset of 73 samples consisted of 3,251,726 fungal sequence reads, with a mean of 44,544 reads per sample (min. 26,494; max. 61,786 reads per sample). These were grouped into 5,276 ASVs, of which 4,472 were taxonomically assigned to the phylum level. Of these, Ascomycota (73.80%) had the most sequence reads, followed by Basidiomycota (12.07%), Mortierellomycota (11.43%) and Chytridiomycota (2.12%). A total of 2,309 ASVs (43.76%, corresponding to 64.09% of reads) could be assigned to at least one guild in the FungalTraits database^1^. Of these, saprotrophs (which include soil saprotrophs, litter saprotrophs, wood saprotrophs, and unspecified saprotrophs) made up 58.34%, plant pathogens 25.82%, and arbuscular mycorrhiza 0.30% of reads.

References of Supplementary Methods S1:

1.	Põlme, S. *et al.* FungalTraits: a user-friendly traits database of fungi and fungus-like stramenopiles. *Fungal Divers.* **105**, 1–16 (2021).
